# Flexible decision-making is related to strategy learning, vicarious trial and error, and medial prefrontal rhythms during spatial set-shifting

**DOI:** 10.1101/2023.12.13.571351

**Authors:** Jesse T. Miles, Ginger L. Mullins, Sheri J. Y. Mizumori

## Abstract

A hallmark of behavioral flexibility is the ability to update behavior in response to changes in context. Most studies tend to rely on error counting around reward contingency or rule switches to measure flexibility, but these measures are difficult to adapt in a way that allows shorter timescale flexibility estimates. Further, choice accuracy does not account for other markers of flexibility, such as the hesitations and decision reversals humans and other animals often exhibit as decisions unfold, a behavior often called vicarious trial and error (VTE). To relate observable information about decision-making to latent aspects like learning and behavioral flexibility, we quantified changes in decision-making strategy using a previously developed, recency-weighted Bayesian inference algorithm. By comparing models of strategy use with decision history to generate strategy likelihood estimates on a trial-by-trial basis, the algorithm enabled us to identify learning points, and served as the basis for the development of a behavioral flexibility score. Aligning flexibility scores to learning points showed that flexibility peaked around estimated learning points and near peaks in VTE rate. However, we occasionally observed VTE during periods of low flexibility, where it often led to incorrect choices, suggesting the likely existence of multiple VTE-types. Additionally, we built on the decades of research suggesting a prominent role for the medial prefrontal cortex in enabling behavioral flexibility by recording field potentials from the medial prefrontal cortex during task performance. We observed changes in different field potential frequency bands that varied with respect to the different behavioral measures we used to characterize learning and decision-making. Overall, we demonstrate the use of multiple measures that jointly assess relationships between learning, behavioral flexibility, and decision-making behaviors. Further, we used these complementary measures to demonstrate that a particular decision-making behavior, VTE, was likely to be a marker of deliberation at some times, and uncertainty at others. Finally, we validate these measures by showing that theta, beta, and gamma rhythms in the medial prefrontal cortex vary with respect to both observable and latent aspects of behavior.

## 1. Introduction

Behavioral flexibility describes the ability to change behavior in response to changing external conditions or internal states (Brown & Tait, 2010; Dalley *et al*., 2004; Hones & Mizumori, 2022; Izquierdo *et al*., 2017; Ragozzino, 2007; Uddin, 2021). Typical tests of behavioral flexibility involve assessing how well subjects perform tasks that require them to update their behavior as task demands change. In humans, one famous example is the Wisconsin Card Sorting Test (WCST), which requires subjects to learn which stimulus quality (color, number, or shape) is rewarded, sort cards based on the currently rewarded quality, and update sorting strategies when rewarded qualities switch (Grant & Berg, 1948; Miyake *et al*., 2000; Uddin, 2021). Similar tasks have been adapted to non-human primates (Butter, 1969; Goudar *et al*., 2023; Mahut, 1971; Moore *et al*., 2005; Roberts *et al*., 1988), and rodents (Becker *et al*., 1981; Izquierdo & Jentsch, 2012; Kolb *et al*., 1974; Ragozzino *et al*., 2003).

In rodents specifically there are two prominent categories of behavioral flexibility tests: reversal learning tasks and set shifting tasks. Reversal learning tasks involve reward contingency changes, typically such that an opposite response or stimulus selection confers reward (*e.g.*, from turn left to turn right). Set shifting tasks, however, require shifts between different task rules that dictate possible reward contingencies (*e.g.*, from turn left to alternate turn directions). Flexibility on either reversal learning or set shifting tasks is often measured by 1) number of trials to a certain performance criterion; 2) number of errors due to use of a strategy that’s no longer rewarded, called perseverative errors; or 3) overall choice accuracy within blocks of trials, sometimes broken down by proximity to switches. All of these measures revolve around distinguishing between possible different types of error and counting the number of errors in different time periods that are dictated by aspects of the task.

Along with the latent, cognitive changes that enable behavioral flexibility, observable behavior is also known to vary. For example, rodents (Muenzinger, 1938; Muenzinger & Gentry, 1931; Tolman, 1926), non-human primates (Kaufman et al., 2015; Medin et al., 1970; Resulaj et al., 2009), and humans (Santos-Pata & Verschure, 2018; Voss & Cohen, 2017) will sometimes appear to pause and/or change the course of a decision as it is carried out, a behavior typically called vicarious trial and error (VTE) but sometimes known as change of mind (Kaufman *et al*., 2015; Resulaj *et al*., 2009). Most initial observations of VTE showed that it tended to happen just before or as rats learned a task (Gentry, 1930; Muenzinger, 1938; Muenzinger & Gentry, 1931; Tolman, 1926). Since then, multiple studies have shown that VTE tends to occur on more difficult decisions (Bett *et al*., 2012; McLaughlin & Redish, 2023; Papale *et al*., 2012, 2016; Schmidt *et al*., 2013), and manipulations that decrease VTE can also impair task performance (Bett *et al*., 2012; Kidder *et al*., 2021; Schmidt *et al*., 2019). This evidence is coherent with the hypothesis that VTE is a marker of deliberative behavior (Redish, 2016) and as such suggests that VTE could serve, along with choice outcome, as another candidate for assessing behavioral flexibility.

Though there is support for the general claim that VTE is associated with behavioral flexibility and deliberation, the relationship between VTE and choice outcome is not always clear or consistent across tasks, and most measures of behavioral flexibility rely on evaluating changes in error rates over multi-trial timescales. While some evidence suggests that VTE and associated behaviors are affected over these longer timescales (George *et al*., 2023; McLaughlin & Redish, 2023; Papale *et al*., 2016), we wanted to measure the association between behavioral flexibility, learning, VTE, and choice outcomes more directly, on a trial-by-trial basis, using behavioral measures that could be calculated independently of one another. To do so, we implemented a spatial set shifting task that required rats to repeatedly switch between blocks of trials where they had to either continually return to the same location (follow a place rule) or alternate between locations on every trial (follow an alternation rule). We utilized a recency-weighted Bayesian inference approach that compares choice history to explicitly modeled behavioral strategies and computes the likelihood that each strategy was being used on every trial (Maggi *et al*., 2023). We identified putative learning points by finding when the target strategy became the most likely strategy for the remainder of the block, and, using changes in likelihoods across trials, we computed a behavioral flexibility score to determine periods of high flexibility that might not be obvious by proximity to learning points or task structure (*e.g.*, block types/switches) alone.

Further, we hypothesized that we should see changes in neural activity if we did indeed identify well-defined measures that meaningfully parsed behavior. Many studies have shown that the rodent medial prefrontal cortex (mPFC) is associated with all of the behavioral measures we were interested in comparing (Durstewitz *et al*., 2010; Euston *et al*., 2012; Euston & McNaughton, 2006; Guise & Shapiro, 2017; Hasz & Redish, 2020a; Hyman *et al*., 2012; Insel & Barnes, 2015; Maggi *et al*., 2018; Powell & Redish, 2016; Pratt & Mizumori, 2001; Rich & Shapiro, 2009). Accordingly, we recorded field potentials from the mPFC during our set shifting task to assess whether activity varied with respect to choice outcome, VTE, learning phase, and flexibility score.

Our results show that learning, behavioral flexibility, VTE, and choice outcomes are typically tightly coupled to one another, but can decouple depending on context. Increases in choice accuracy, VTE rates, and flexibility scores aligned to identified learning points. In support of the often-claimed role for VTE in deliberation, VTE trials were more likely to end in correct choices, and correct VTE trials were more likely to have a higher flexibility score. However, VTE on trials during low flexibility periods were more likely to lead to errors, and incorrect VTE trials were more likely to have lower flexibility scores, suggesting that VTE may sometimes be a marker of uncertainty, not deliberation.

Additionally, both observable and latent behavioral measures were associated with changes to power distributions in different mPFC field potential frequency bands. Specifically, trials with VTE showed elevated theta and beta compared to non-VTE trials, and periods when learned strategies could be exploited after learning were associated with stronger gamma. Taken together, these results suggest that the confluence of these behavioral measures can be used to delineate behavioral contexts, as exemplified by our demonstration that VTEs can be separated as either deliberative or uncertain. Moreover, we strengthen our behavioral findings by showing variations in mPFC activity that track both observable and latent behavioral measures. Overall, these results help link learning, behavioral flexibility, variations in decision-making behaviors, and changes in mPFC physiology through mutually corroborative evidence.

## 2. Results

We utilized a spatial set shifting task (**Figure 1A**) that required rats to either continually return to the same location (use a place rule) or alternate between locations on successive trials (use an alternation rule). This design is similar to Meyer-Mueller *et al*., (2020), except we use a plus-maze instead of T-maze, with randomly chosen start arms to ensure that only strictly allocentric, place strategies (as opposed to egocentric, body-turn strategies) will be successful. Because prior results show differences in VTE rates for egocentric compared to allocentric navigation (Schmidt *et al*., 2013), but no consistent changes during switches between different egocentric strategies (Meyer-Mueller *et al*., 2020), we analyzed whether there were performance differences in allocentric place compared to allocentric alternation strategies.

**Figure 1.**
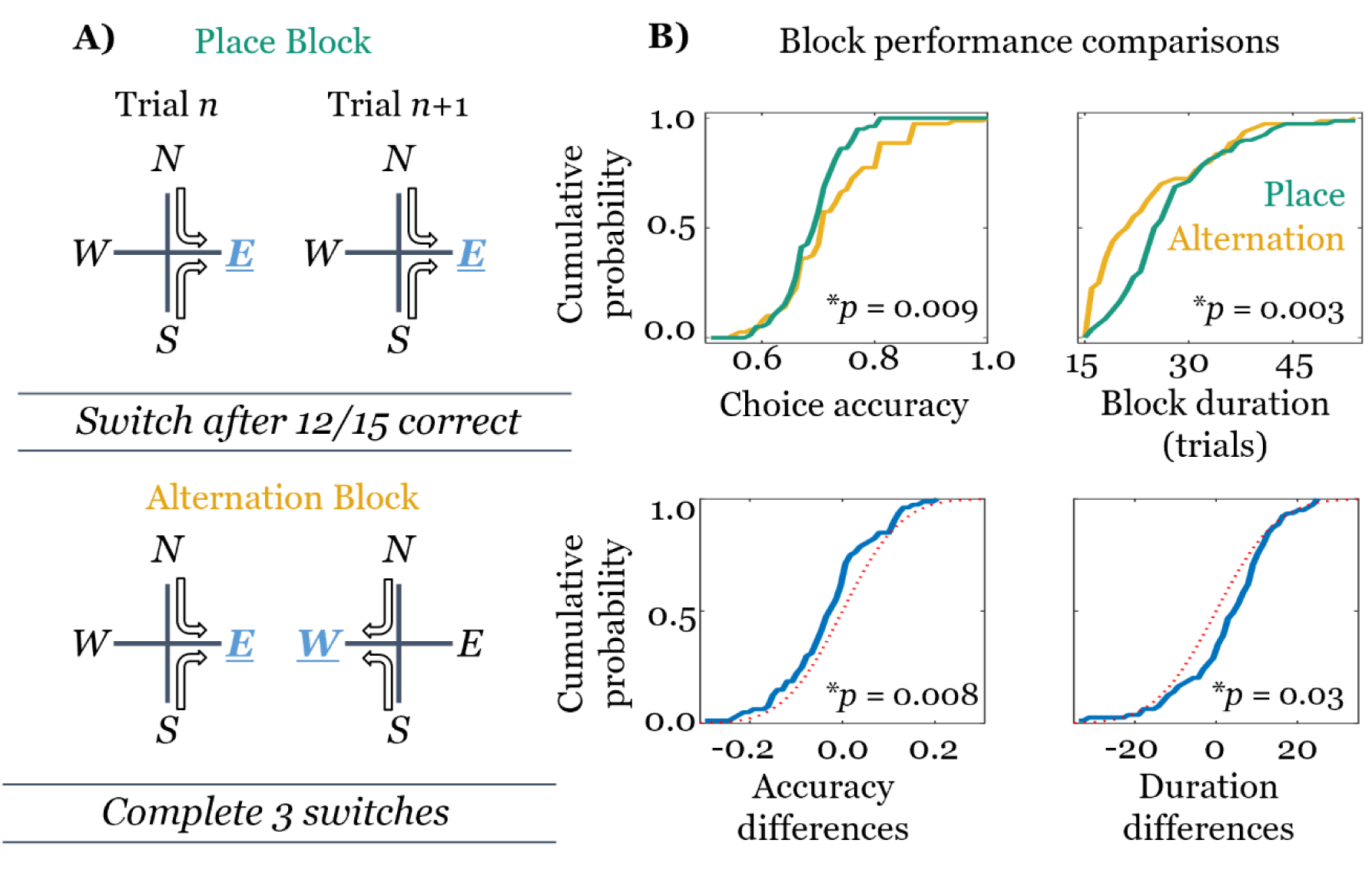
Performance differs according to task rule. **A)** Diagram of the task rules. Place blocks (top) reward continual visits to a particular (E or W) arm, and alternation blocks (bottom) reward alternation between E and W blocks on successive trials. Sessions consist of 3 block switches. Switches occur when 12 of the previous 15 choices were consistent with the target rule. **B)** Multiple measures of performance are better for alternation blocks compared to place blocks. Unpaired cumulative distributions for choice accuracy within a block are leftward shifted for place blocks (top left, green line), while alternation block durations are leftward shifted (top right, gold line). *p*-values are calculated using unpaired, two-sample, two-sided T-tests. Within-session, paired comparisons suggest the same conclusion (bottom). Differences between choice accuracy in adjacent place and alternation blocks (bottom left solid blue line, place minus alternation) are shifted to the left of zero, while differences in block duration are shifted to the right of zero (bottom right solid blue line, place minus alternation). p-values are calculated using signed rank tests. Red dashed lines are zero-mean (left) or median (right), standard deviation-matched normal cumulative distributions for comparison.

Choice accuracy distributions for alternation blocks were right-shifted compared to place blocks (**Figure 1B**, top right; 2-sample, 2-tailed, T-test; t = 2.66; 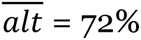, 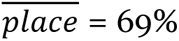; *p =* 0.009; *d* = 0.42; overbars represent the sample mean). Block duration distributions were non-normally distributed for alternation trials, which showed cumulative probability bunched around the lower block duration limit (15 trials). This distribution was left-shifted compared place block durations (**Figure 1B**, top right; 2-sample, 2-tailed Wilcoxon rank-sum test; Z = −2.94; *alt_m_* = 23.3, *place_m_ =* 25.8, where subscript *m* denotes median; *p =* 0.003; *d* = −0.3). Within session comparison of the adjacent pairs of place and alternation blocks (**Figure 1B**, bottom right) yields the same result for choice accuracy difference (t = 2.71; 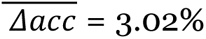; *p* = 0.008; *d* = 0.30) and block duration difference (t = 2.71; 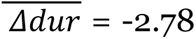; *p* = 0.03; *d* = −0.24). While these differences are consistent, note that they are not large (∼3% performance difference and ∼3 trial duration difference; Cohen’s *d* values below 0.5).

We implemented a previously developed algorithm that uses recency weighted Bayesian inference to identify changes in strategy learning (Maggi *et al*., 2023). Comparing decision history (not choice outcome) to modeled strategies allowed us to identify learning by finding when a rat’s most likely strategy matched the target task rule (see **Figure 2A** for an example of the estimation output). Conceptually, we regard the learning point as splitting a block into a pre-learning point exploratory period, where different strategies are tested, and an exploitation period, where memory can be used to guide decision-making. Since the learning point was identified without explicit reference to choice outcome, seeing increases in the likelihood of a correct choice with respect to the putative learning point would corroborate that it had been correctly identified. As expected, average choice accuracy aligned to learning points showed a striking increase just prior to the learning point, remaining elevated for several trials after. As shown in **Figure 2B**, average choice accuracy (dashed horizontal line) for the 15 trials up to the learning point (dashed vertical line) is 63.4% and the lower bound of an estimated 95% confidence interval exceeds that value starting 1 trial before the learning point, peaks 1 trial after the learning point, and remains above the pre-learning point average for 7 trials after the learning point (data shown for *n* = 40 sessions, where each point within 15 trials on either side of the learning point is the average choice accuracy across 4 blocks; though we see the same result using *n* = 13 subjects with averages across 4 to 24 blocks).

**Figure 2.**
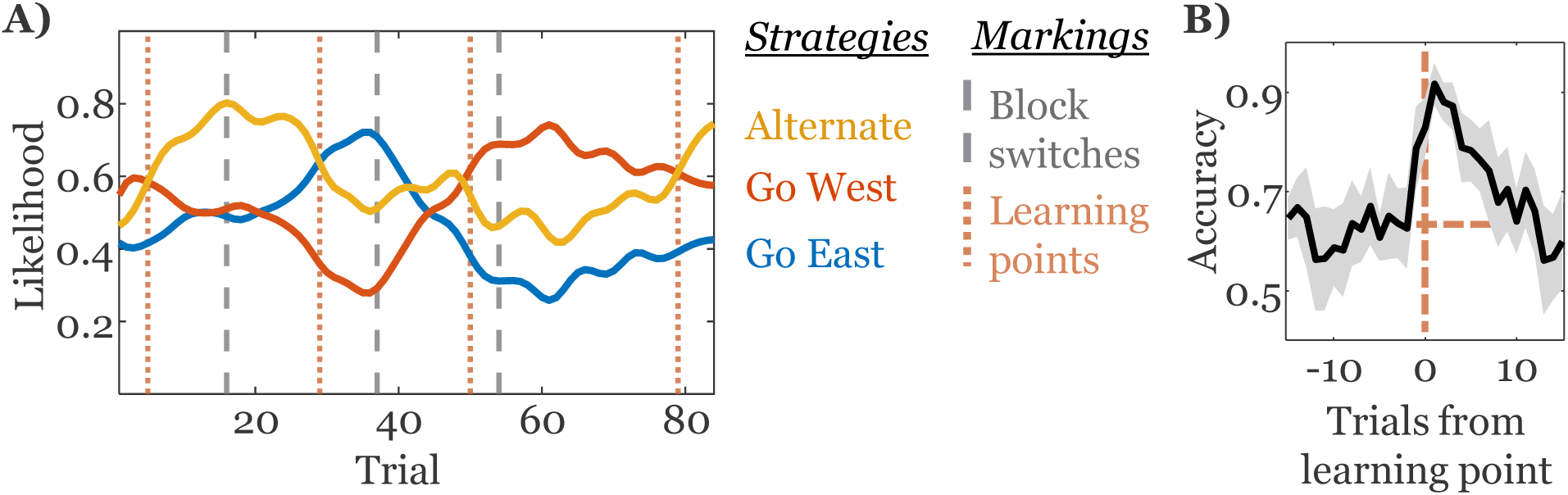
Identifying learning points from strategy likelihoods. **A)** We modeled three strategies – go east (blue), go west (orange) and alternate (yellow). Learning points are indicated with vertical, orange, dotted lines, and block switches are indicated by vertical, grey, dashed lines. The average choice accuracy aligned to learning points is shown in **B)**. A shaded 95% confidence interval surrounds the average. The horizontal dashed line indicates the pre learning point average, the vertical dashed line indicates the learning point.

As mentioned, prior reports show VTE rate differences for different types of strategy (Schmidt *et al*., 2013). In our task, both strategies had right-skewed, overlapping probability distributions of VTE rates (**Figure 3A**, top right; 2-sample, 2-tailed Wilcoxon rank-sum test; Z = 0.36; *alt_m_*= 15%, *place_m_ =* 14%; *p =* 0.72; *d* = 0.03). Other studies suggest that the relationship between VTE and choice outcome is, if nothing else, task dependent, so we calculated the within-session difference between the number of VTE on trials with correct and incorrect choices. For this task, VTE was far more likely on correct choice trials. In fact, there were only 4 sessions (10%) where VTE led to errors more often than correct choices, and out of those 4, VTE was far more likely to precede errors in only 1 (**Figure 3B**; t = 5.04; 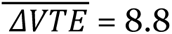; *p* < 0.001; *d* = 0.80, gold lines denote sessions with at least as many VTEs leading to errors).

**Figure 3.**
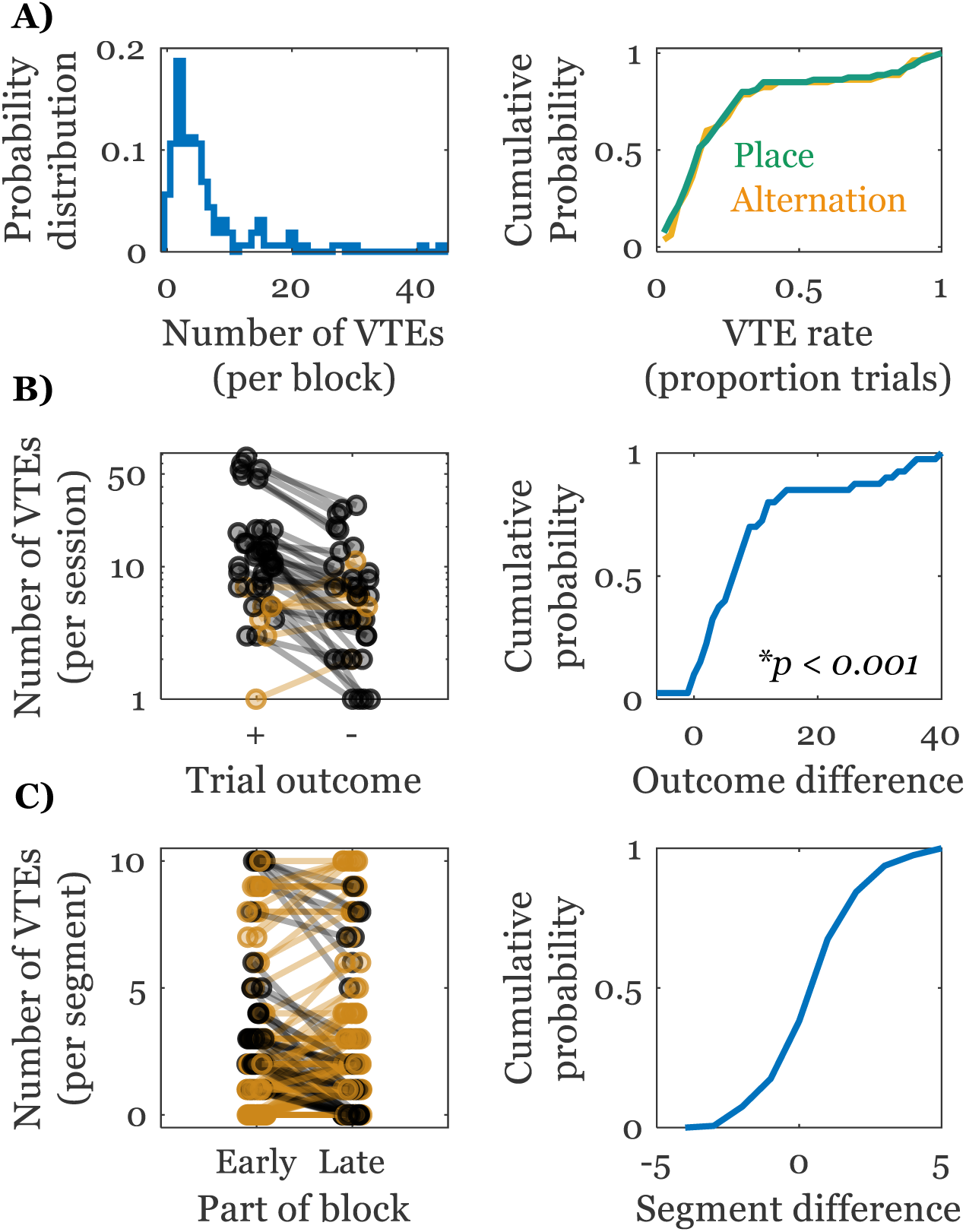
Summary of VTEs by session and block. **A)** The left histogram shows the number of VTE trials per block, while the right shows proportions of VTEs per block. The right column shows that there are no differences in how proportions of VTE per block are distributed for either place (green) or alternation (gold) blocks. **B)** Looking at within session differences in VTE that led to correct compared to incorrect choices shows that VTE was far more likely to lead to a correct choice. Raw numbers are shown on the left (with log y-axis on left, not log x on right), and the distribution of within session differences are on the right. **C)** Although VTE is more likely to lead to correct choices, there are no differences in the probabilities of VTEs occurring early compared to late in blocks.

The current and historical literature don’t seem to have come to a consensus on how VTEs should unfold throughout the course of learning. Some report that VTE in navigation or location-based tasks decrease over time as learning occurs (Jackson, 1943; Kemble & Beckman, 1970) in a task dependent manner (Goss & Wischner, 1956), but in other tasks VTE has been shown to stay elevated throughout, supposedly depending on the task difficulty (Gentry, 1930; Tolman, 1948). Our task ensures that the current contingency has been learned at the end of a block but is unknown at the beginning of a block. Thus, we asked if there were differences in the number of VTE trials in the first 10 trials of a block compared to the last 10. We saw that there were no differences in the number of VTE early vs late in the block (**Figure 3C**; t = −0.43; 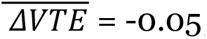; *p* = 0.67; *d* = −0.03, gold lines denote blocks with at least as many VTEs later in the block).

Because VTE has been suggested to track task learning and has been proposed as a behavioral marker for deliberation, we asked whether changes in VTE rates align to learning points. Indeed, **Figure 4C** shows that there are significantly elevated VTE rates from 3 trials before to 1 trial after the learning point (see **Methods** for estimation of VTE rates and statistical analysis paradigm), further validating the association between learning and VTE without appealing to choice accuracy.

**Figure 4.**
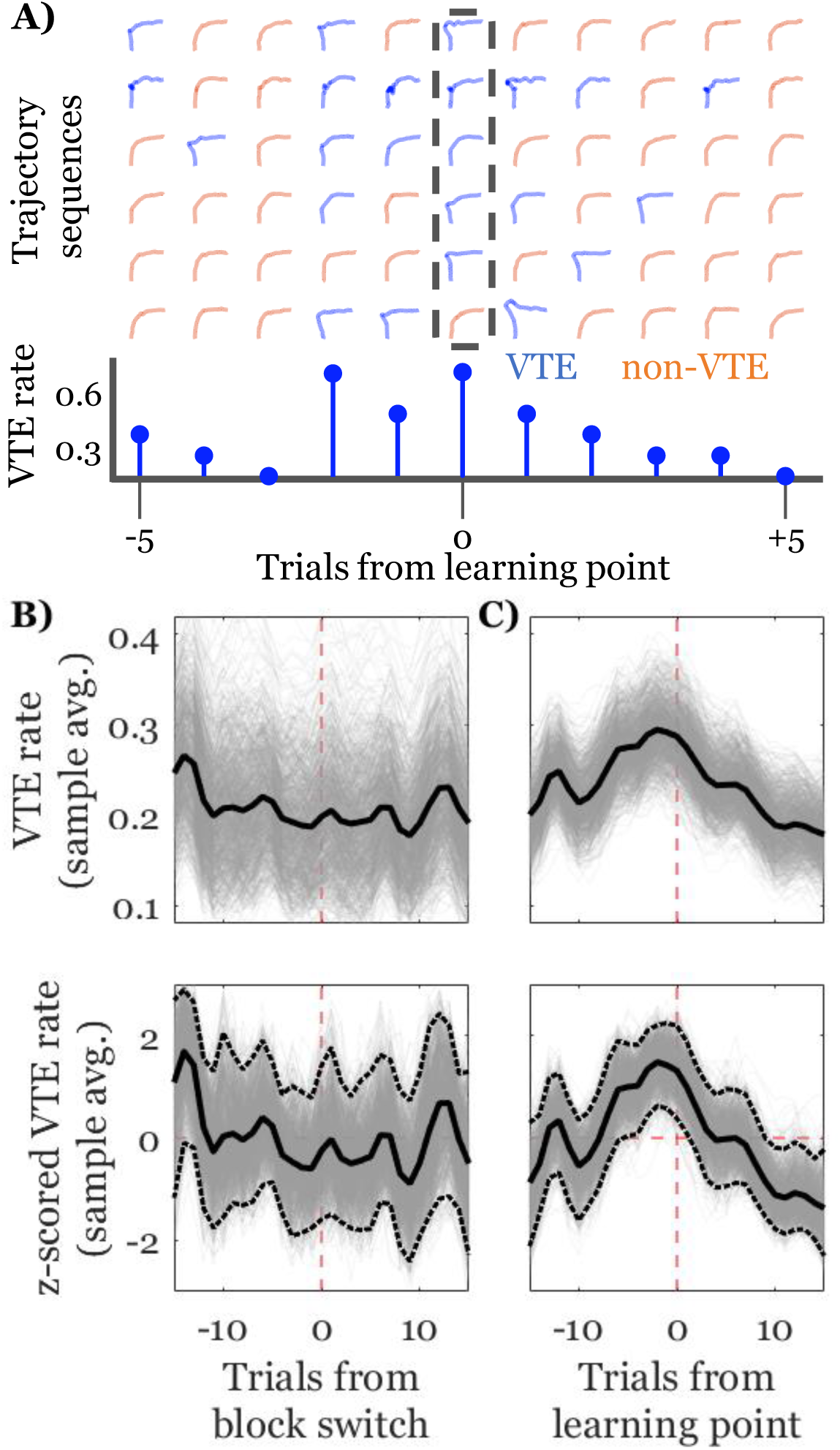
VTE rate changes align with learning points. Each row in **A)** shows trajectories from sequences of trials aligned to estimated learning points (dashed rectangle/trial 0 on the bottom axis) for a given block. Blue trajectories have been identified as VTEs and orange trajectories have been identified as non-VTEs. The stem plot shows VTE rates for this sample of six sequences. Note the general increased rate for trials near the learning point. The top panels in **B)** and **C)** show changes in VTE rate with respect to block switches on the left and estimated learning points on the right (dashed, vertical, red lines). For all plots, bold, black lines are the averages of 1000 iterations of a hierarchical bootstrap to estimate VTE rate timeseries from binary vectors; light grey lines are the individual iteration results. Top panels show raw rates and bottom plots show z-scored rates. Dashed, horizontal, red lines in the lower plots show the mean for comparison. Dotted black lines show the 2.5 and 97.5 percentiles of the distribution. Dotted black points on the graph above zero show trials where less than 2.5% of estimates across iterations were below zero for that trial (trials −5 to 1), and vice-versa for points below zero (trial 9 to 15).

These elevations contrast with VTE rates aligned to block switches, which hover around the average for almost the entire window (**Figure 4B**).

Changes in strategy likelihoods suggest updates in choice behavior, allowing us to define a behavioral flexibility measure based on trial-by-trial strategy likelihood changes (**Figure 5A**). Importantly, this method allowed us to measure behavioral flexibility without reference to choice outcome, which could mask instances where subjects did switch between strategies, but neither strategy matched the rule they were meant to follow. Further, it allowed us to assess flexibility trial-by-trial instead of over periods of trials. Much like choice accuracy and VTE rate dynamics, aligning sequences of flexibility scores to the learning points showed that there were consistent increases in flexibility starting just before the learning point that end several trials after (**Figure 5B**). Note that although learning points and flexibility scores are defined by strategy likelihoods, flexibility scores can and do vary – sometimes dramatically – away from the learning point. Likewise, sometimes flexibility scores were lower during learning points when transitions happened slowly. Thus, this result was expected, but not guaranteed.

**Figure 5.**
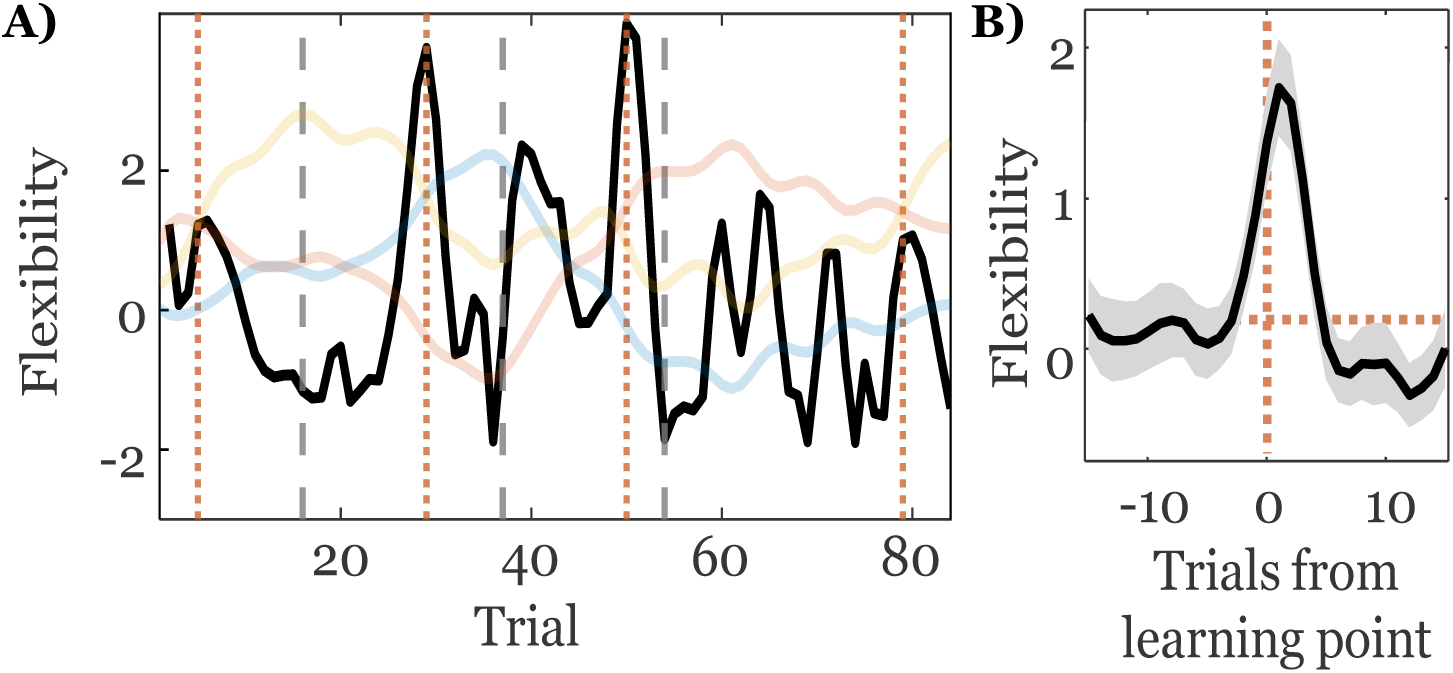
Flexibility score example and relation to learning point. **A)** Bold, black line indicates flexibility scores for one example session (same as Figure 2A, shown in background) Learning points are indicated with vertical, orange, dotted lines, and block switches are indicated by vertical, grey, dashed lines. The average flexibility score aligned to learning points is shown in **B)**. A shaded 95% confidence interval surrounds the average. The horizontal dashed line shows the pre-learning point average, the vertical dashed line shows the learning point.

The associations between VTE rates, flexibility dynamics, and choice accuracy changes provide strong support for the claim that VTE can serve as a marker for deliberation. However, not all VTE occurred around learning points, some VTE occurred during periods of low flexibility, and some VTE led to errors. Thus, we asked whether there may have been another, non-deliberative type of VTE. We know that VTE near learning points happens as choice accuracy and flexibility are high, but to see if opposing relationships existed as well, we asked if incorrect VTEs were associated with lower flexibility scores, and if VTEs that happened in inflexible periods were more likely to be incorrect. Indeed, incorrect VTEs had significantly leftward shifted flexibility scores (**Figure 6A**; t = 9.40; 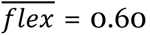, 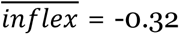; *p <* 0.0001; *d* = 0.57). We defined a set of criteria that determined whether a VTE occurred during a flexible or inflexible period. First, any trial within two trials of a learning point was considered flexible (regardless of flexibility score). Second, a trial had to be more than three trials prior to the end of a block (unless it was within 2 trials from the learning point). Third, any trial with a flexibility score in the top 60^th^ percentile was considered flexible (unless it was within three trials from a block switch). Similarly, inflexible periods could not be within two trials of the learning point (regardless of flexibility score) and had to have flexibility scores in the bottom 40^th^ percentile (see **Methods** for further descriptions). Within-session, paired comparisons showed that choice accuracy was significantly higher during flexible, learning related VTE than VTE during inflexible periods that didn’t border the learning point (**Figure 6B**; t = 5.23; 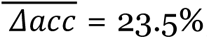; *p <* 0.0001; *d* = 1.01). Together, these data suggest that VTE can happen in at least two contexts – one that suggests a deliberative process, and another which looks more like uncertainty.

**Figure 6.**
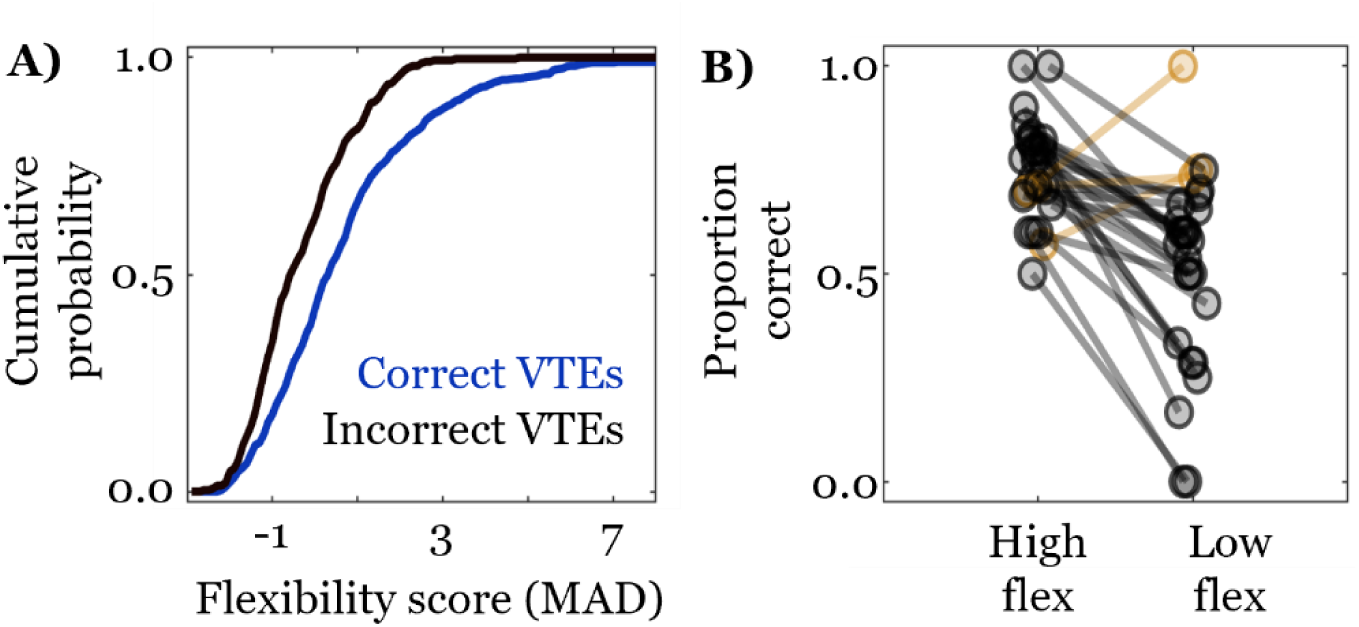
Evidence for multiple VTE types. **A)** Correct VTEs are more likely to have higher flexibility scores than incorrect VTEs. VTEs in the top 40^th^ percentile of flexibility scores are more likely to be correct than VTEs in the bottom 40^th^ percentile.

The medial prefrontal cortex has been repeatedly implicated in strategy switching tasks and tasks that require spatial working memory. As such, we recorded mPFC field potential rhythms from a subset of rats used in the behavioral dataset (*n* = 3) to see if mPFC rhythms tracked any aspects of behavioral context formation that we were able to define (see **Figure 7** for approximate recording locations and field potential examples). We examined three mPFC rhythms, each suggested to have a role in linking cognition to behavior in rodents; 1) the theta rhythm (6 – 12 Hz), which tends to synchronize with hippocampal theta when spatial working memory is taxed (Benchenane *et al*., 2010; de Mooij-van Malsen *et al*., 2023; Hallock *et al*., 2016; Hyman *et al*., 2010; Jones & Wilson, 2005; Negrón-Oyarzo *et al*., 2018; Stout *et al*., 2023; Tavares & Tort, 2022); 2) the beta rhythm (15 – 30 Hz), which tends to be present during decision-making (de Mooij-van Malsen *et al*., 2023; Jayachandran *et al*., 2023; Symanski *et al*., 2022); and 3) the gamma rhythm (40 – 100 Hz), which has been associated with working memory, learning, and sensory information processing (Cansler *et al*., 2022; de Mooij-van Malsen *et al*., 2023; Negrón-Oyarzo *et al*., 2018).

**Figure 7.**
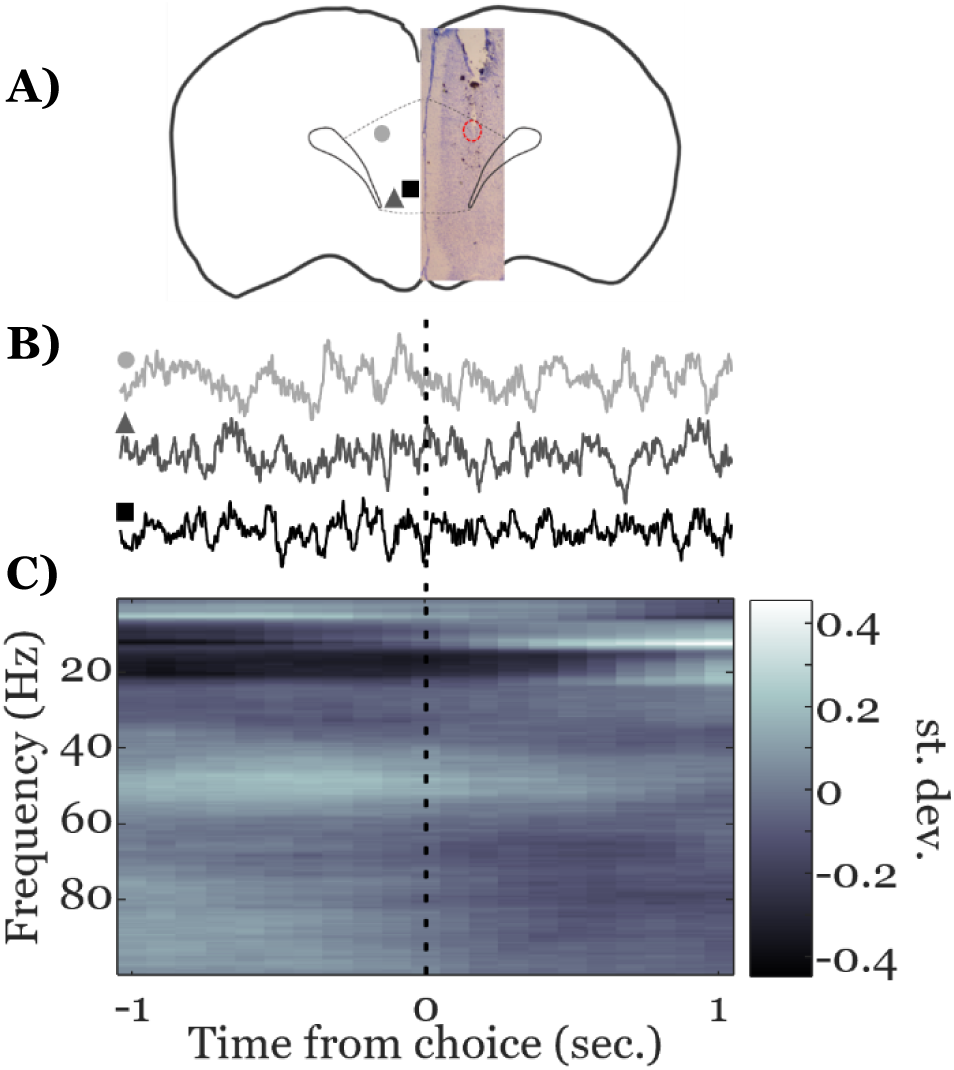
Histological placement and LFP examples. **A)** Recording sites were spread throughout the prelimbic cortex. Different shapes denote tip locations for the three different animals in the dataset. Sites span approximately +3.7 to +4.3 mm anterior to bregma. **B)** LFP examples from each of the three recording sites, marked by corresponding shapes and shades from **A)**. Each recording is aligned so the center of the trace is when the rat crosses the center of the platform, with 1 second before and 1 second after on either side. **C)** Average spectrogram of all LFPs from across choices. Each trial’s spectrogram is mean subtracted and scaled by the standard deviation, across time, for every frequency.

As shown in **Figure 7A**, the three recording sites we analyzed were located in anterior prelimbic cortex (see *Electrophysiology* section of **Methods** for approximate coordinates). Field potentials were aligned to decision points (**Figure 7B**), converted into time-frequency representations, and normalized by trial across a 6 second window containing 3 seconds before and 3 seconds after the decision point (average across all trials shown in **Figure 7C**). On average, most of the variance in the spectrogram appears distributed within the *a priori* defined frequency bands described above.

We tested whether mPFC field potential rhythms were related to any of the behavioral measurements used to define context by comparing rhythms on trials with opposing contextual components. There were four main components used to delineate behavioral contexts: choice outcome, whether VTE occurred, flexibility, and learning phase. To make paired, within session comparisons, we employed a similar hierarchical bootstrap sampling technique used to generate VTE rate curves. First, subjects were sampled, then, for each subject, a random sample of sessions were drawn, and, within each session, we computed the mean difference in the strength of rhythmic activity for the different frequency bands on opposing trial types (*i.e.*, correct minus incorrect, VTE minus non-VTE, exploit phase minus explore phase, and high flexibility minus low flexibility). We refer to the first element in the pair (*e.g.*, correct trials) as the condition trial and its opposite (*e.g.*, incorrect trials) as the comparison trial. Repeatedly sampling in this way produces a posterior distribution of differences. We assume that if there were no difference between condition and comparison trials, distributions should be centered at zero with a roughly even proportion of the data on either size of the mean. As such, we quantified the strength of evidence for a particular rhythm varying with respect to a given contextual component by the probability that its distribution sat above zero. If none of the data for a given distribution were above zero, the probability value (*P*) would be zero, and this would be very strong evidence that those condition trials had weaker activity than their accompanying comparison trials in that frequency band. At the other extreme, if all of the distribution was above zero, this would be a probability value of 1, and strong evidence that the condition had stronger rhythmic activity in that band than the comparison.

Results for different opposing trial combinations, separated by rhythm, are shown in **Figure 8**. Shades of the distributions vary such that darker shades indicate stronger evidence that condition trials have weaker rhythms than comparison trials for trial type, whereas lighter shades indicate stronger evidence of that rhythm’s presence on condition trials than comparison trials. An additional measure, analogous to Cohen’s D for one sample distributions, is reported in the upper corner for each distribution. The value’s magnitude measures how many standard deviations the distribution’s mean is from zero, and its sign tells in which direction. The first row, comparing correct and incorrect trials, shows that both theta and gamma distributions are close to zero, with little indication that trial types differ. Gamma, however, appears to be more consistently weaker on correct trials (**Figure 8A**, Gamma, *P* = 0.21, *d* = −0.84). Interestingly, all rhythms tend to have stronger increases during VTE trials, with very strong evidence for beta (**Figure 8B**, Beta, *P =* 0.98, *d* = 2.36) and strong evidence for theta (**Figure 8B**, Theta, *P* = 0.93, *d* = 1.49) on VTE trials. Only gamma appears to show any difference in the post-learning, exploit phase compared to the explore phase, with strong evidence for higher gamma during the exploit period (**Figure 8C**, Gamma, *P* = 0.95, *d* = 1.62). Comparing high and low flexibility trials shows weak evidence that theta may be higher on high flexibility trials while gamma is more consistently lower on high flexibility trials (**Figure 8D**, Gamma, *P* = 0.18, *d* = −0.85).

**Figure 8.**
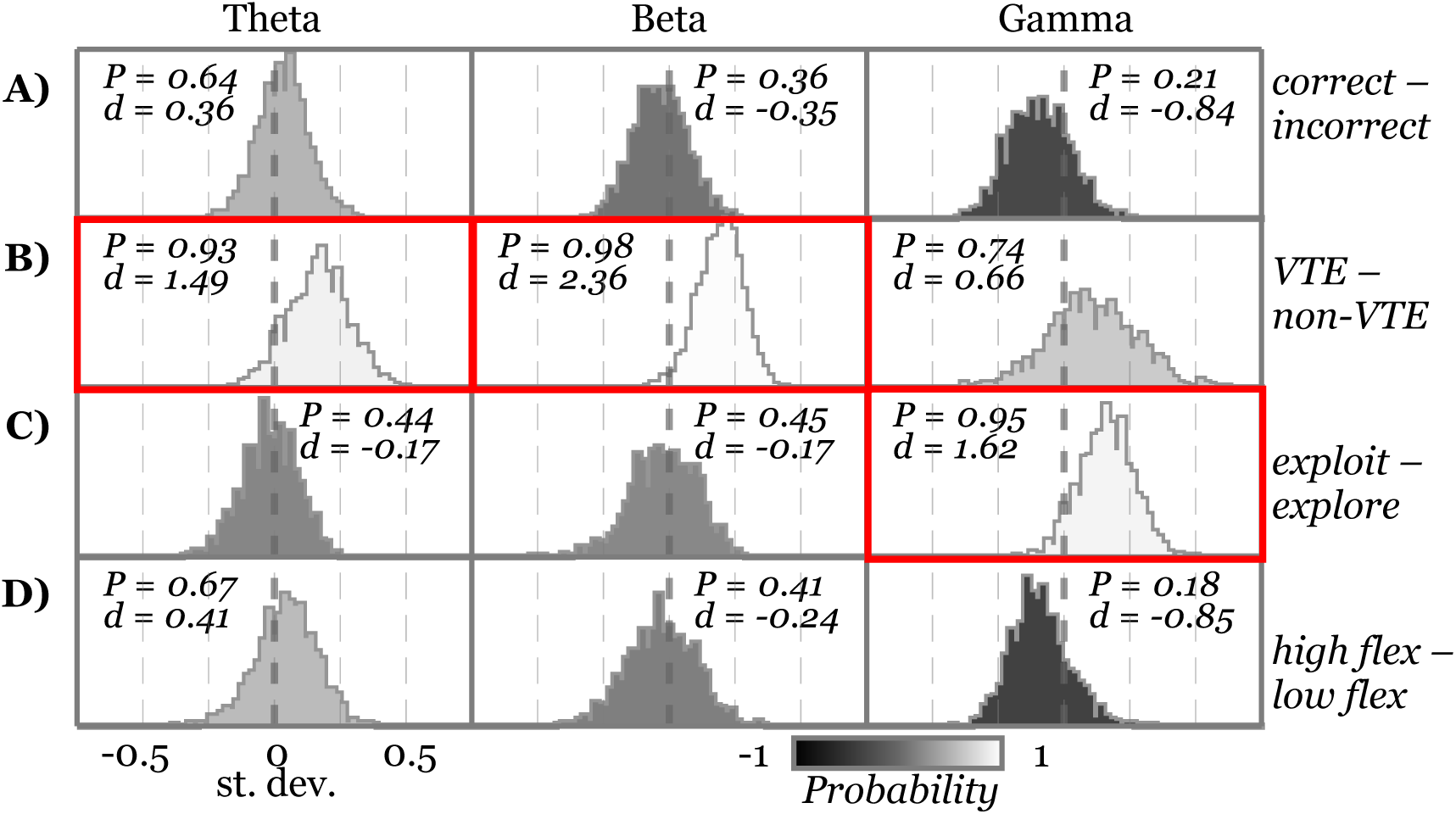
mPFC rhythms vary based on contextual components. Each box in the grid shows the distribution of differences for hierarchically sampled trial comparisons. The type of trial comparison is shown on the left side of the figure, outside the grid of distributions. The left column shows comparisons for the theta rhythm, the middle for the beta rhythm, and the right for the gamma rhythm. The probability of the distribution falling above zero is indicated by *P* in one of the upper corners of each plot, and a one sample analog to Cohen’s D is indicated by *d* underneath. Each distribution is shaded according to its *Probability* value, as shown by the gradient below the grid. Distributions outlined in red have *P* ≥ 0.9 and *d* ≥ 1.0, and are considered to represent strong evidence for a difference between groups.

## 3 Discussion

Behavioral flexibility is a complex phenomenon that could manifest in many different ways, but our typical understanding of it primarily focuses on a single measure – choice outcome. VTE behavior has been documented for nearly a century, but it has been difficult to reconcile descriptions of its function. This study sought to supplement our understanding of behavioral flexibility and fill in some of the gaps in the VTE literature by analyzing VTE with respect to other streams of behavioral data that also occurred on a trial-by-trial basis during a dynamic decision-making task. To do so, we estimated strategy likelihoods from rule-based models, which enabled learning point identification, and developed a behavioral flexibility measure based on changes in strategy likelihood estimates. We show that choice accuracy, VTEs, and flexibility scores all increased surrounding learning points. Further, we show that VTEs were far more likely to be correct than incorrect in this task, and that correct VTEs were more likely to occur on trials with higher flexibility scores, suggesting a typical role in deliberation. However, we also found VTE that occurred during periods of low flexibility and were often wrong, indicating that VTE may sometimes reflect uncertainty instead of deliberation. Finally, we showed that these behavioral measures often had distinctive relationships to mPFC field potential activity. Covariations between mPFC activity and specific behavioral measures provide strong validation that our analyses effectively partitioned behavior into relevant, naturalistic epochs.

As mentioned, VTE-like behaviors are present in humans (Iggena *et al*., 2023; Santos-Pata & Verschure, 2018; Voss & Cohen, 2017), while inflexible, perseverative, behavior and difficulty with executive control are common measures in clinical diagnoses of neurological disorders (Uddin, 2021). Because any behavior that can be identified and modeled based on decision-making is amenable to the analysis workflow we’ve utilized, and many decision-making tasks require some active trajectory toward a decision, we hope that the framework described here will be of general interest to behavioral neuroscientists asking both basic and clinical questions.

### Expanding analyses of behavioral flexibility

Based on the simple premise that behavioral flexibility manifests as changes in strategy use, we were able to score flexibility on a trial-by-trial basis and track its changes with respect to task dynamics. We verified that these scores did indeed track flexibility by showing their strong alignment with putative learning points, increases in choice accuracy, and peaks in VTE rate, which have also been proposed as a marker of flexible, deliberative behavior. Having a continuous scale that enables trial-by-trial identification of high and low flexibility based on statistically derived cutoffs can be useful for providing additional context to other behavioral measures, as we show in **Figure 6**. In our case, extra context about flexibility showed that VTE on low flexibility trials were likely to result in errors. By putting these facts together, we concluded that VTE resulting in error on low flexibility trials was likely to represent uncertainty about the decision, which differs from the typical interpretation of VTE as a deliberative behavior.

Another benefit of having flexibility scores that do not depend on choice outcome is the ability to identify periods of high flexibility but low choice accuracy. This occurs, for example, when a subject switches from a prior strategy to a new strategy that does not match the target. In our task this would happen if the prior strategy was *go east* and the current strategy is *go west*, but a subject started alternating instead of switching immediately to *go west*. Identifying these periods could prove particularly useful for trying to disentangle learning, reward processing, or attention from flexibility. Increased flexibility after block switches, but longer exploration phases, for example, could indicate that flexibility is not affected directly, but something about subjects’ ability to stabilize behavior is impaired. On the other hand, unaffected exploration periods coupled with long post-learning periods could indicate that subjects struggle to exploit their newly learned strategies. Both of these are distinct from a situation in which flexibility remained low after block switches due to continual elevated likelihood of a prior strategy, which would indicate perseveration.

Though our analysis of mPFC activity did not reveal strong changes in rhythmic activity as a function of flexibility, it did suggest that gamma rhythms may be weaker in the high flexibility state (**Figure 8d**, right column). Interestingly, when combined with our other observations, we might expect incorrect, inflexible trials during the exploitation period to have some of the strongest gamma activity. These are trials where a rat chose incorrectly, despite a recent history of correct responses. Importantly, because we’ve shown that learning phase and gamma strength are related, we might expect a similar choice history pattern during the exploration phase to have weaker gamma band activity. These distinctions highlight the utility of contextualizing behavior when interpreting neural data.

### Reconciling VTE findings and contextualizing behavior

There are several hypotheses about what VTE is and why it happens. Most of the initial reports claimed that VTE tended to happen just before or as rats learned a task (Gentry, 1930; Muenzinger, 1938; Muenzinger & Gentry, 1931; Tolman, 1926). Muenzinger (1938) and Gentry (1930) noted that, although they had assumed the pause and reorient behavior we now call VTE would primarily reflect active sampling of sensory stimuli, rats still showed VTE when sensory environments were the same and thus not useful in determining where to go for reward. This suggested that the behavior was not simply used to compare sensory information but may instead indicate comparison of past and present experience. Additionally, rats continued to VTE throughout their learning and training during difficult tasks, but they would typically stop after they had learned to consistently make simple sensory discriminations, often interpreted as having formed a habit (Gentry, 1930; Muenzinger, 1938; Tolman, 1948).

In the context of more recent experiments, the repeated observation that VTE tends to occur just after new reward contingency or rule switch and decrease farther into blocks (Blumenthal *et al*., 2011; Kidder *et al*., 2023; Meyer-Mueller *et al*., 2020), are in accordance with the early observations that VTE is linked to learning. The majority of modern research treats VTE as a marker of deliberation, but inconsistency in how VTE relates to choice outcome (George *et al*., 2023; Kidder *et al*., 2021; Meyer-Mueller *et al*., 2020; Schmidt *et al*., 2013), suggests that what VTE represents or is used for may not have a unitary explanation (Goss & Wischner, 1956). This possibility was reported in Gentry (1930), who showed that some rats seemed to VTE consistently while never learning proficiently, while others performed exceptionally well, but did not exhibit the typical decline in VTE rates. Her characterization was that VTE consistently associated with poor performance could indicate never having truly learned the task, while VTE during high performance marked continued deliberation. Tolman similarly claimed that VTE during difficult sensory discriminations may persist because comparison and indecision persist, while its increase during initial learning on easy sensory discriminations is because rats concurrently learned which sensory stimuli (visual/auditory) to associate with reward, as well the discriminative reward contingency (black vs white/toward tone vs away from tone) itself (Tolman, 1948).

Separating VTE into subtypes based on context may help explain some of the idiosyncrasies in how different studies have reported on and conceptualized VTE. For example, silencing the nucleus reuniens has been shown to increase VTE during inflexible periods of perseverative responding, leading to incorrect choices (Stout *et al*., 2022). In our framework, we would interpret VTE in this context as indicative of uncertainty instead of deliberation. Similarly, Schmidt *et al*., (2013) reported that VTE typically led to an error and was most likely on difficult trials – in other words, during times when uncertainty was likely high. However, they also reported VTE during periods of high task proficiency and *after* errors, times when animals could deliberate based on prior experience and understanding. These situations and our demonstration that VTE could be separated based on outcome and flexibility score suggest that future work should take steps to determine whether VTE reflects a deliberative behavior or one driven by uncertainty.

### Neural manipulations and VTE

Several mPFC (Kidder *et al*., 2021, 2023; McLaughlin & Redish, 2023; Schmidt *et al*., 2019) and hippocampus (Bett *et al*., 2012, 2015; Blumenthal *et al*., 2011; Hu & Amsel, 1995; Meyer-Mueller *et al*., 2020) manipulation studies seem to agree that manipulating these regions is unlikely to increase VTE rates. Curiously, manipulating structures that connect these regions (*e.g.* the amygdala, perirhinal cortex, and nucleus reuniens) does increase VTE (Kemble & Beckman, 1970; Kreher *et al*., 2019; Stout *et al*., 2022). One hypothesis for why this may be is that both the hippocampus and mPFC are specifically involved in *enabling* deliberative VTE behavior. When one of these structures isn’t functioning as usual, the deliberation process may fail altogether. Mechanistically, it may be that sequential hippocampal activity generates an array of possible options (Johnson & Redish, 2007; Kay *et al*., 2020) while the mPFC evaluates those options prior to choices (Hasz & Redish, 2020b; Redish, 2016; Schmidt *et al*., 2019; Tang *et al*., 2021; Zielinski *et al*., 2019). If either the representation of possibilities in the hippocampus or evaluative process in the mPFC fails to occur, VTE may become less likely to happen. In contrast, if both processes proceed as they normally would locally, but are unable to coordinate correctly because of interruptions in connecting circuitry, VTE may be just as likely – if not more likely to occur – but related to uncertainty instead of deliberation.

### Neural activity in the mPFC

Intriguingly, not only are our behavioral measures self-consistent and useful for defining latent behavioral contexts, they are also consistent with neural measures of mPFC activity. Our analysis of mPFC field potentials shows that different contextual components are associated with different activity states. The primary goal of these analyses was to provide further validation for our methods of parceling behavior and the measurements we used. Nevertheless, our results provide some insights into how mPFC rhythms reorganize with respect to behavior. As an example, two of the strongest relationships we see are higher mPFC theta and beta on VTE trials compared to non-VTE trials (**Figure 8**, 2^nd^ row, left and middle columns). This is in line with results showing hippocampal activity changes during VTE (Amemiya & Redish, 2018; Johnson & Redish, 2007; Miles *et al*., 2021; Papale *et al*., 2016; Schmidt *et al*., 2019) and the evidence for hippocampal-prefrontal interactions during VTE (Hasz & Redish, 2020b; Schmidt *et al*., 2019; Stout *et al*., 2022). The beta rhythm, specifically, has recently been shown to synchronize the mPFC and hippocampus via brief activity bursts in the nucleus reuniens during an odor sequence memory task (Jayachandran *et al*., 2023), and our result provides further evidence that beta-rhythmic activity in the mPFC component of this tri-partite circuit is crucial for memory-guided decision-making.

Much like we treat VTE as a binary event – it either happens or does not – trials can either be correct or not. Despite the strong evidence for broad spectral changes in the mPFC on VTE trials compared to non-VTE trials, we do not see particularly strong evidence of differences in any band based on trial outcome. There is perhaps some hint that gamma is weaker on correct compared to incorrect trials, but it could be that latent factors play a larger role in modulating mPFC rhythms relative to choice accuracy.

The broadest latent context change we define – the shift from strategy exploration to exploitation after the learning point – is more clearly accompanied by changes in gamma rhythms (**Figure 8**, 3^rd^ row, right column). This is in line with prior work in rodents showing that prefrontal explore-exploit relationships are impaired if the mPFC is inactivated (Birrell & Brown, 2000; Laskowski *et al*., 2016; Ragozzino *et al*., 1999, 2003), as well as studies of mPFC units in rats showing that both individual units (Jung *et al*., 1998; Rich & Shapiro, 2009) and population level encodings (Guise & Shapiro, 2017; Hasz & Redish, 2020a; Maggi & Humphries, 2022; Malagon-Vina *et al*., 2018) track strategy switches. An alternative explanation could be that subjects were more likely to be correct during exploitation periods, and gamma during choices was related to expected choice outcome. If this were the case, we would expect to see the same pattern of increased gamma-rhythmic activity on correct compared to incorrect choices, but, as mentioned, we see a trend in the opposite direction (**Figure 8**, 1^st^ row, right column). This suggests that it is not expected outcome driving the difference in gamma fluctuations during explore and exploit trials. In support of these observations, recent work in humans with intracranial EEG recordings has shown that there are changes in gamma band activity during the transition between exploration and exploitation (Domenech *et al*., 2020).

Another latent context measure is flexibility magnitude, which, according to our data, may also be associated with changes in the gamma rhythm (**Figure 8**, 4^th^ row, right column). When split into high and low flexibility trials, gamma has a very similar distribution to gamma differences related to trial outcome. These two measures could very well be related, as both flexibility and choice accuracy peak around learning points, but there is not as clear a relationship between flexibility and learning phase. Flexibility can be quite variable during exploration as different strategies are tested (*e.g.,* around trials 40 and 60 in **Figure 5A**), and flexibility also typically transitions quickly from high at the beginning of the exploit period to low within several trials (**Figure 5B**). While all of these measures likely have some individual relationship to gamma, it’s also likely that those relationships are not independent of one another, and it remains to be seen how they covary. Still, our results suggest that both latent and explicit behavioral measures appear to have distinct associations with gamma in the prefrontal cortex.

### Conclusion

This study characterized learning, decision-making behaviors, and mPFC activity during a spatial set shifting task. By quantifying changes in strategy use, we proposed a new way of calculating behavioral flexibility and show that flexibility scores aligned with increases in decision-making accuracy and VTE-rates that accompanied learning. At other times, relationships between these patterns broke down. Examining trials with atypical behavioral patterns enabled reinterpretation of similar looking behaviors as cognitively distinct. Finally, we showed that these measures, particularly VTE and learning phase, show distinctive relationships to mPFC rhythms.

## 4 Methods

### Subjects, apparatus, training protocol, and behavioral task

Food restricted (85% of body weight) Long Evans rats (n = 13, Charles River Laboratories) were trained to perform a spatial set shifting task. Sessions were run on an elevated plus maze (black plexiglass arms, 58 cm long x 5.5 cm wide, elevated 80 cm from floor), with moveable arms and reward feeders controlled by custom LabView 2016 software (National Instruments, Austin, TX, USA) that tracked the rat and automatically raised and lowered arms based on positions recorded by a SONY USB web camera (Sony Corporation, Minato, Tokyo) acquiring frames at approximately 35 Hz. Rats were initially habituated to handlers and the maze for however long it took them to comfortably interact with handlers, forage for pellets on the maze, and habituate to maze movement and noise (several days to one week).

Task training started with a forced choice paradigm, where rats learned the trial structure, consisting of leaving its pseudo-randomly chosen “North” or “South” starting arm, then navigating to an “East” or “West” arm for a 45 mg sucrose pellet reward (TestDiet, Richmond, IN, USA). Initially, rats completed 5 to 7 trials that forced them to either alternate between reward sites or return repeatedly to the same East or West location, regardless of their starting point. This number was increased until rats could do 10 trials of each reward contingency in under 45 minutes, at which point they were given a free choice version of the same task.

In the free-choice training version, we did not change start arms on error trials until rats made a certain number of correct choices, and initially started with a low choice accuracy criterion for switching between blocks (typically around a 70% success rate in an 8 to 10 trial window). As rats started completing all reward contingencies (alternate, go east, go west), we increased the criterion for success and minimum number of trials per block, and decreased the number of correct trials needed before errors no longer influenced start arm switches. This procedure was tailored to each rat until it could complete 4 reward contingencies (two alternation blocks and two place blocks; one East, one West) in less than 150 trials, with start arms pseudo-randomly chosen for all trials. We also ensured that two alternation blocks did not occur back-to-back.

Testing sessions followed the same trial structure as the final training sessions. As mentioned above, start arms were “pseudo-random”. This was done to ensure that between 50 – 60% of trials within 15 trial stretches had start arm switches. Doing so increased the number of switches compared to what you’d expect from random draws, while slightly decreasing the number of long (4 to 10 trial) sequences where the start arm stayed the same, and eliminating sequences without a switch that were longer than that. For this dataset, all rats completed three switches, though not all completed the 4^th^ block. A total of 40 sessions from 13 rats were analyzed with all but one rat contributing at least two sessions.

### Position tracking and VTE identification

We identified VTEs in much the same way as Kidder *et al*. (2023). Briefly, we took the videos that tracked coarse body location during the task and used DeepLabCut (DLC) version 2.2 (Mathis *et al*., 2018; Nath *et al*., 2019) to identify the rats’ heads. We started with the same model trained in Kidder *et al*. (2023) and retrained a new iteration with additional labeled data from the set shifting experiments. Each training attempt used NVIDIA GEFORCE GTX 1080 GPU with 500,000 iterations. Trajectories with vicarious trial and error (VTE) were detected by projecting the position data into principal component (PC) space and clustering the PC-representations of the trajectories with hierarchical agglomerative clustering. Before projection, all trajectories were aligned and standardized to the same starting and ending positions and interpolated (or linearly subsampled, if necessary) to have the same number of points. Visual inspection of the clustering in PC space naturally formed two clouds in low dimensional plots, and distance-based dendrograms cut to give two clusters separated trajectories with VTE from non-VTE trajectories, though with some errors.

To mitigate the errors, we reassigned certain trajectories initially not identified as VTE but with x-positions that crossed a certain threshold into the VTE category, and used the combination of a low *z-ln(idphi)* measure (Bett *et al*., 2012; Blumenthal *et al*., 2011; McLaughlin & Redish, 2023; Papale *et al*., 2012; Schmidt *et al*., 2013, 2019; Stout *et al*., 2022) and a failure to cross lower x-position boundary to reassign any VTEs that may have been mistakenly identified. Informal inspections of randomly sampled data subsets after classification suggest that this method is between 80% and 90% accurate, which is in-line with supervised classification methods and near the threshold for interrater agreement (Miles *et al*., 2021).

### Estimating strategy likelihoods, learning points, and flexibility scores

We adopted the procedure from Maggi *et al*. (2023) to estimate explicitly modeled strategy likelihoods, trial-by-trial, based on choice history and a recency-weighted decay factor to account for the inherent non-stationarity in behavior associated with strategy switching. Minor changes to strategy templates allowed us to model allocentric versions of the strategies instead of the egocentric versions originally used; we simply added a column to the data processing input that parsed “East” and “West” choices instead of “Left” and “Right” (though the algorithm can easily process both types of reference frame if both types of input are given). As in the original report, we used 0.9 as the parameter controlling the strength of the recency weighting. One addition we made was to add several randomly permuted trials to the beginning of sessions prior to processing with the algorithm. This dampened some of the algorithm’s initial large swings in likelihood estimates that were due to limited trial history. Further, we smoothed likelihood timeseries with a 5-trial Gaussian window to increase the reliability of learning point identification. As suggested in the original paper, we identified learning points as the trial when the target strategy became the most likely.

Our rationale for calculating flexibility scores from changes in strategy likelihood is based on the notion that decision-making patterns shifting to become consistent with a different strategy is the sign of flexible behavior in a set shifting task. Thus, for each trial, the flexibility score is the absolute difference in strategy likelihoods from trial t-1 to trial t, summed across strategies. For each session, this value is normalized by median absolute deviation (robust Z-score) because values tend to deviate more dramatically in the positive than negative direction, but the results remain the same when a normal, standard deviation-based Z-score is used.

Flexible periods, used for testing whether there were multiple VTE types, were defined based on three criteria. First, trials on either side of the learning point were automatically considered flexible, regardless of their flexibility score. Trials that were three trials before the end of a block were automatically *not* considered flexible, unless they were within one trial of the learning point. Third, any remaining trials in the top 60% of flexibility scores were considered flexible, while any in the bottom 40% of flexibility scores were not considered flexible. When analyzing choice accuracy on VTE trials during flexible compared to inflexible periods, we excluded sessions where there were three or fewer trials of either type. Results for this analysis did not change if we changed the ratios of flexible and inflexible VTEs that could be included (*e.g*., based the third criterion on flexibility score instead of percentile), but this often changed the proportion of data we were able to use.

### Statistical quantification

Critical values were set at 0.05. When sample sizes were large, and distributions looked nearly normal, two-tailed T-tests were used. If making within session comparisons, one-sample tests of differences were done, comparing the empirical distribution to what would be expected for a distribution with 0 mean. When distributions were strongly skewed, we performed two-tailed Wilcoxon rank sum tests if the data were not paired, and two-tailed Wilcoxon signed rank tests when they were paired. Learning point aligned choice accuracy and flexibility score data were not subjected to formal statistical testing – instead, 95% confidence intervals were estimated as if they were from a normal distribution (using Z-values). These curves are presented as averages and confidence intervals across sessions (n = 40), but looked nearly identical when comparing across subjects (n = 13).

Learning point aligned VTE “rasters” and a peri-learn point VTE rate averages suggested that VTEs likely aligned to learning as well, but to test this we used hierarchical bootstrapping (Saravanan *et al*., 2020). This helped safeguard results from bias introduced by uneven data collection between subjects while also acknowledging that multiple measurements from individuals were not independent. It also allowed us to create rate distributions out of binary data. Subjects were sampled randomly with replacement enough times to match the total number of subjects in the dataset (n = 13), then, for each subject, a certain number of blocks was also randomly selected with replacement (8 for learning point and 6 for block switch aligned sequences of VTE data – equivalent to two sessions of data). Average VTE rates were calculated for these samples and smoothed with a 5 trial Gaussian window across trials. Distributions were formed by repeating this procedure 1000 times. Significance was determined by asking which trials had more than 97.5% of their (Z-scored) iterations above 0. Results were tested without smoothing, using larger smoothing windows, with different numbers of iterations, using median absolute deviations instead of standard deviation Z-scoring, using shorter and longer sequences of trials, and across many random seeds, all leading to the same conclusion. The only thing that sometimes changed was the number of trials surrounding the learning point that show significantly elevated VTE rates.

### Electrophysiology

A subset of the rats (n = 3) had neural recordings from the mPFC. Recordings were collected from custom-built tetrode micro-drives with Intan headstages and Open-Ephys acquisition systems, as described in (Kidder *et al*., 2021; Miles *et al*., 2021). All mPFC recordings were localized to the prelimbic cortex (**Figure 7A**), between approximately 3.7 and 4.3 mm anterior to bregma.

Recording windows analyzed were determined by finding the choice point, identified as the point closest to the center of the platform for each trajectory, and then extending four seconds before and after that point. For each window, time-frequency spectrograms were calculated using Chronux’s `mtspecgramc` function (Bokil *et al*., 2010), using a two second window, 100 msec overlap, 7 tapers, and a bandpass window of 0.5 to 100 Hz, generating a time-frequency matrix for each decision. After converting power values to decibels, we Z-scored each frequency component along the time dimension for each trial to normalize the data (see **Figure 7C** to see the average representation of this matrix). Since we were working with *a priori* frequency bands and were only interested in decision-based activity, we averaged activity across each frequency band from −1 second before to 1 second after the decision point, meaning each trial gave a single average value for theta, beta, and gamma band activity.

We analyzed field potential activity in relation to different behavioral measures by creating hierarchically sampled bootstrap distributions (Saravanan *et al*., 2020) for specified pairs of conditions (**Figure 8**). Each measure had a reference trial type, which we called the condition trial, and an opposing trial type called the comparison trial. For example, trials could either be correct (a condition trial) or incorrect (a comparison trial). This allowed us to perform within-session, paired comparisons for each pair by subtracting the activity averaged across all sampled trials in the comparison group from the condition group. For each hierarchical sample, we used 3 individuals, 4 sessions, and 40 trials from each group (condition and comparison), all sampled with replacement. Thus, each difference is the average of activity from the 40 trials from the condition group minus the average activity of the 40 trials from the comparison group, and a member of the overall distribution is the mean of these differences across the 12 sessions drawn for that sample iteration. This was repeated 1000 times to generate the distributions shown in **Figure 8**.

## Acknowledgements and contributions

We would like to thank Arman Khan and Trinity Charles for their help with training, data collection, and troubleshooting the paradigm, and Dr. David Gire, Dr. Kevan Kidder, Victoria Hones, and Maeve Bottoms for helpful comments throughout the study. This research was supported by National Institute of Mental Health research grant MH119391 to Dr. Sheri Mizumori. The authors do not declare any conflicts of interest.

JTM implemented the paradigm, collected data, performed formal analyses, wrote/adapted software, and wrote the original draft of the manuscript. GLM collected data and helped design and troubleshoot the paradigm. SJYM supervised the project, acquired funding, and contributed revisions to the manuscript.

